# Genetic characterization of a recombinant myxoma virus leap into the Iberian hare (*Lepus granatensis*)

**DOI:** 10.1101/624338

**Authors:** Ana Águeda-Pinto, Ana Lemos de Matos, Mário Abrantes, Simona Kraberger, Maria A. Risalde, Christian Gortázar, Grant McFadden, Arvind Varsani, Pedro J. Esteves

## Abstract

Myxomatosis is a lethal disease of wild European and domestic rabbits (*Oryctolagus cuniculus*) caused by a Myxoma virus (MYXV) infection, a leporipoxvirus that is found naturally in some *Sylvilagus* rabbit species in South America and California. The introduction of MYXV in the early 1950s into feral European rabbit populations in Australia and Europe demonstrate the best documented field example of host-virus coevolution following a cross-species transmission. Recently, a new cross-species jump of MYXV has been suggested in both Great Britain and Spain, where European brown hares (*Lepus europaeus*) and Iberian hares (*Lepus granatensis*) were found dead with lesions consistent with those observed in myxomatosis. To investigate the possibility of a new cross-species transmission event by MYXV, tissue samples collected from a wild Iberian hare found dead in Spain (Toledo region) were analyzed and deep sequenced. Our results report a new MYXV strain (MYXV Toledo) in the tissues of this species. The genome of this new strain encodes three disrupted genes (*M009L*, *M036L* and *M152R*) and a novel 2.8 KB recombinant region that resulted from an insertion of four novel poxviral genes towards the 5’ end of its genome. From the open reading frames inserted into the MYXV Toledo strain, a new orthologue of a poxvirus host range gene family member was identified which is related to the MYXV gene *M064R*. Overall, we confirmed the identity of a new MYXV strain in Iberian hares that we hypothesize was able to more effectively counteract the host defenses in hares and start an infectious process in this new host.

## Introduction

Myxoma virus (MYXV), a poxvirus belonging to the *Leporipoxvirus* genus, is the etiological agent of myxomatosis which is a highly lethal viral disease of wild and domestic European rabbits (*Oryctolagus cuniculus*) [1]. The classical form of the disease is characterized by systemic spread of the virus, overwhelming the immune system, and the development of secondary skin lesions called ‘myxomas’ [2, 3]. Mortality rate varies between 20-100%, according to the grade of virulence of the MYXV strain [3]. The virus has its natural host in the South American tapeti, or forest rabbit (*Sylvilagus brasiliensis*), where it causes an innocuous and localized cutaneous fibroma at the inoculation site [2]. Related poxviruses to MYXV are found in other *Sylvilagus* species in North America: the Californian MYXV strains, for which the natural host is *Sylvilagus bachmani* (brush rabbit), and rabbit fibroma virus (RFV) found in *Sylvilagus floridanus* (eastern cottontail) [2, 4]. MYXV not appearing to cause significant clinical disease in the natural *Sylvilagus* hosts, though being highly pathogenic to the naive *Oryctolagus* host, made it a classic example of a pathogen that is highly virulent in a new host species with no evolutionary history of adaptation to that pathogen.

In 1950, with the urge of controlling the infesting population of European rabbits in Australia, a MYXV strain originally isolated in Brazil (standard laboratory strain [SLS]) was used as a biological agent [1]. The release in France in 1952 of a different Brazilian isolate of MYXV (Lausanne [Lu] strain) resulted in the establishment and spread of MYXV in Europe, including the United Kingdom (UK) [5]. After an initial massive reduction of the wild rabbit populations (>99%) in both Continents, a substantial decline in the case fatality rates occurred as a result of natural selection for slightly attenuated viruses, but also due to an increased resistance to myxomatosis in the rabbit populations [4, 6, 7]. It has been recently shown that the convergent phenotype of viral resistance observed in Australia, France and UK rabbit populations was followed by a strong pattern of parallel evolution, a consequence of selection acting on standing genetic variation that was present in the ancestral rabbit populations in continental Europe [8].

The susceptibility of other leporids species to MYXV has been tested in controlled experiments, while evidence of myxomatosis in wild leporid populations have been seldom reported. Using a California MYXV strain four different North American *Sylvilagus* species (*S. audubonii*, *S. floridanus*, *S. idahoensis* [now *Brachylagus idahoensis*] and *S. nuttallii*) developed tumors following mosquito transfers, but these failed to be mosquito-infective lesions [9]. Three of these *Sylvilagus* species (*S. audubonii*, *S. floridanus* and *S. nuttallii*) when infected with the Brazilian Lu strain also developed prominent tumors, however this time the South American strain produced mosquito-infective lesions [10]. On the other hand, black-tailed jackrabbits (*Lepus californicus*) inoculated with Californian MYXV did not form tumors [9]. In wild populations of European hare (*Lepus europaeus*) cases of myxomatosis have been reported sporadically and in small number. In the past, the confirmation of the disease arose from injecting rabbits with tissues from dead hares and replicating its typical clinical symptoms [11]. Most recently, in 2014, for the first time a case of myxomatosis in a European brown hare in Great Britain was confirmed using electron microscopy and a PCR of a skin lesion [12].

Recently, in late summer-fall of 2018, the first cases of myxomatosis in Spanish wild Iberian hare (*Lepus granatensis*) populations were reported, mainly in the Andalusia and Castilla-La Mancha regions. The Spanish Ministry of Agriculture, Fisheries and Food, and the Institute for Game and Wildlife Research identified what appeared to be a cross-species transmission into a new leporid species. Iberian hares were found in moribund state, with signs of blindness, weakness and disorientation, and consequently analyzed in different laboratories. Here, using culturing and deep sequencing, we genetically characterize for the first time a recombinant MYXV isolated from an Iberian hare carcass exhibiting classical symptoms of myxomatosis collected in Toledo province, Spain during the 2018 outbreak (referred to as MYXV Toledo).

## Methods

### Sampling and pathology

An adult Iberian hare (*L. granatensis*) female, was found dead on 21st of August 2018 in La Villa de Don Fabrique municipality in the Toledo province of Spain. The hare manifested lesions compatible with myxomatosis in European rabbits (Figure 1) and was completely emaciated (kidney fat index = 0). On arrival at the laboratory, duplicate samples (4mm diameter) were taken from eyelid, ear and vulva and stored in RNAlater and without preservative, at −80ºC. For the histopathological study, representative samples of the main organs and tissues, were fixed in 10% buffered formalin for 48-72 hours at 22±2°C, and then, dehydrated in a graded series of ethanol, immersed in xylol, and embedded in paraffin wax using an automatic processor. Sections were cut at 4 µm and stained with hematoxylin and eosin (H&E), following standard procedures.

**Figure 1:**
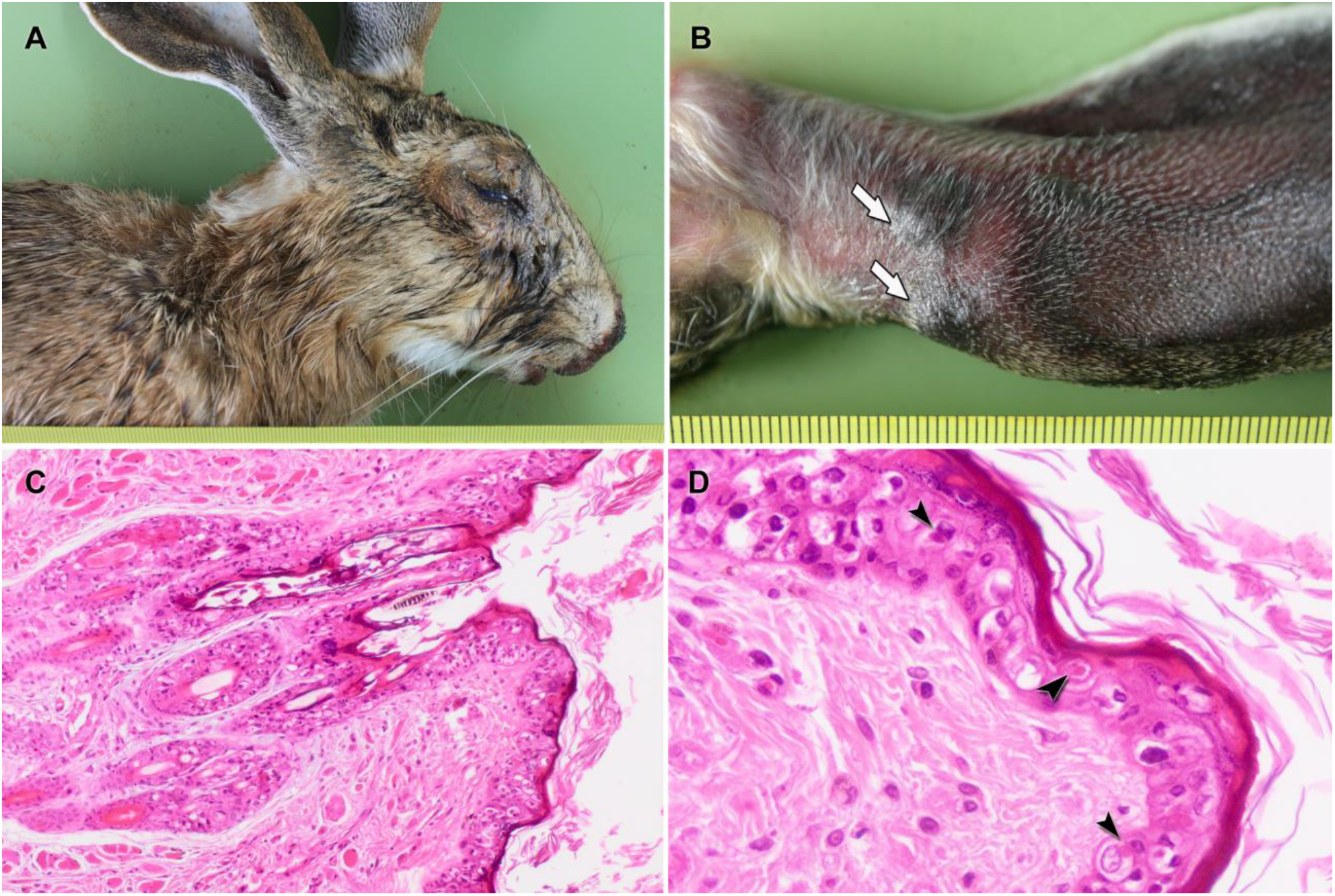
Iberian hare with myxomatosis-compatible lesions. A) Blepharitis and conjunctivitis with seropurulent discharge. B) Myxomas at the base of the left ear (arrows). C) Severe acanthosis of the eyelid skin, with hyperkeratosis. D) Ballooning degeneration of the epidermal cells, and intracytoplasmic eosinophilic inclusion bodies in the eyelid skin (arrowheads).

### Cell lines

European rabbit RK13 kidney epithelial cells (Millipore Sigma, USA) were maintained in Dulbecco’s modified Eagle medium (HyClone, USA) supplemented with 10% fetal bovine serum (FBS), 2 mM L-glutamine, and 100 U/ml of penicillin/streptomycin. Cells were maintained at 37°C in a humidified 5% CO_2_ incubator.

### Isolation, amplification and purification of the new Myxoma virus (MYXV Toledo) strain

Samples from lesions of the eyelid and urogenital regions of an Iberian hare (*L. granatensis*) specimen were manually homogenized. A small volume (5-10 μl) of the processed tissues was used to inoculate confluent RK13 cells monolayers in a 6-well plate and allowed to incubate at 37°C. At 2 days after infection, distinctive MYXV foci were visualized using a Leica DMI6000 B inverted microscope. To proceed with the virus isolation, infected cells were harvested, freeze-thawed at −80°C and 37°C for three times and sonicated for one minute to release the viruses from infected cells. The virus was inoculated back onto a confluent RK13 cells monolayer in a 150 mm dish and incubated at 37°C for 48 hours. Cells were collected to perform a serial dilution and the one with the best individualized foci (dilution 10^−5^) was used for inoculating a new 150 mm dish. After 2 days of infection, a last round of cell harvest, freeze-thaw cycles and sonication was done before proceeding to virus amplification into twenty 150 mm dishes. Purification of the virus through a 36% sucrose cushion was performed as described before [13]. Titration of the number of replicating infectious units of virus was determined by crystal violet foci staining of the infected RK13 cell monolayers, while the total number of viral particles was counted using the NanoSight NS300 instrument (Malvern Panalytical, USA).

### Viral nucleic acid extraction, Illumina sequencing and *de novo* assembly of the genome

Total viral nucleic acid was extracted from 200 µl of the viral prep using a phenol-chloroform extraction protocol as previously described [14]. The viral DNA was used to generate a 2×100 bp Illumina sequencing library and this was sequenced on a Illumina HiSeq4000 (Illumina, USA) at Macrogen Inc. (Korea). The paired-end raw reads (40,730,938 reads) were *de novo* assembled using metaSPAdes v3.12.0 [15] with kmer of 33, 55 and 77. The *de novo* assembled contigs then assembled into a genome length contigs using MYXV-Lu (GenBank accession # MK836424) as a scaffold, primarily to resolve the terminals redundancy. The quality of the final assembly was verified by mapping the raw reads back to the genome using BBMap [16].

### Genome analysis

All MYXV and poxvirus RefSeqs were downloaded from GenBank on the April 9, 2019. Global alignments of the MYXV with the genome determined in this study were carried out using MAFFT [17]. ORFs in the genome were determined with ORFfinder (https://www.ncbi.nlm.nih.gov/orffinder/) coupled with a local MYXV ORF database generated from the MYXV genomes. ORFs that did not have any similarity to MYXV ORFs were analyzed using BLASTn and BLASTx sequence queries [18]. All pairwise identities (nucleotide and protein) were calculated using SDV v1.2 [19].

Protein sequence alignments of the newly derived poxvirus virion protein, thymidine kinase, host range protein and poly(A) Polymerase subunit were used to inferred maximum likelihood phylogenetic trees using PHYML 3.0 [20] with substitution models JTT+G, WAG+G+F, JTT+G+F and JTT+G+F respectively, determined using ProtTest [21]. Branches with aLRT support of <0.8 were collapsed using TreeGraph2 [22].

## Results and Discussion

The natural host for MYXV is the South American tapeti (South American strains) [2, 4]. As expected from predictions of long-term virus/host co-evolution, MYXV strains are highly adapted to their natural hosts, causing only benign cutaneous fibromas [4]. However, when another susceptible host becomes available to the virus transmission system, in this case the European rabbit (*Oryctolagus cuniculis*), a successful cross-species transmission can occur. Indeed, when MYXV first entered the European rabbit host, it was immediately pathogenic and caused close to 100% mortality. After the use of MYXV in the 1950s to control feral rabbit populations in Australia and Europe, rapid co-evolutionary changes occurred in both rabbit host and virus, due to increased resistance of rabbit populations and the appearance of less virulent virus strains [8, 23]. In 2014, a study reported the presence of a myxomatosis-like disease in the European brown hare (*Lepus europaeus*) [12]. However, a MYXV strain capable of infecting hares has not been previously genetically characterized. More recently, reports of abnormal mortalities in Iberian hares were described in the Spanish regions of Andalucía, Castilla-La Mancha, Extremadura, Madrid and Murcia. The animals found in the hunting grounds presented with inflammation of the eyelids, conjunctivitis and also inflammation of the perianal area, symptoms consistent with classic rabbit myxomatosis.

In this study, a new MYXV strain (MYXV Toledo) was isolated and sequenced from an Iberian hare found in Toledo province (Figure 1A, E) that presented the classical lesions of myxomatosis, including a bilateral blepharitis and conjunctivitis, and a swollen vulvar and anal region. The basal third of the left ear presented two myxoma-like lesions of 5 mm diameter (Figure 1B).Moreover, epistaxis and strong congestion of the trachea were observed, whereas the lung was swollen and presented few petechial hemorrhages. Histopathology analysis of the eyelid skin revealed the typical proliferation and ballooning degeneration of the epidermal cells, containing single large rare, intracytoplasmic, round and eosinophilic inclusion bodies (Figure 1C). In this tissue, a severe acanthosis with erosion and ulceration was observed. Blepharoconjunctivitis lesions were also associated with an inflammatory cell response in the underlying dermis, with infiltration of large macrophage-like cells, diffuse edema and fibrin deposition (Figure 1D). In the lung, mild congestion, alveolar edema and hemorrhages were observed. These vascular lesions were also recorded in the liver and kidneys.

### Comparison of Lausanne strain with the newly discovered Toledo MYXV strain

From the collected samples, a new MYXV strain was isolated which we have named MYXV Toledo strain (MYXV-To). The *de novo* assembled genome is 164,579 bps. This genome was aligned to MYXV-Lu strain (GenBank accession # AF170726.2) for preliminary analysis. The MYXV-To genome (GenBank accession # MK836424) was found to be ~2,800 bp longer than the one reported for the MYXV-Lu strain (161,777 bp) [24]. Based on the published genome sequence, the MYXV-Lu strain has a total of 171 genes (12 of which are duplicated in the TIRs regions) [3, 24] and can be divided into three regions: the terminals, 14.1 kb extending from the left TIR (*M0005.1L* to *M011L*) and 23.1 kb extending from the right TIR (*M143R* to *M000.5L*) mostly contain genes involved in the MYXV virulence and host subversion, while the central 124.5 kb region (*M012L* to *M142R*) includes a mixture of virulence genes and essential viral genes conserved across all poxviruses [2, 25]. Of the 159 different MYXV-Lu strain encoded gene products, all were ~99% identical to those of the MYXV-To strain, with the exception the ORFs *M009L*, *M152R* and *M036L*. Furthermore, we identified a novel insertion of ~ 2,800 bp within the *M009L* gene that spans the 12,236 to the 15,082-bp region of the left end of the MYXV-To genome (Figure 2).

**Figure 2:**
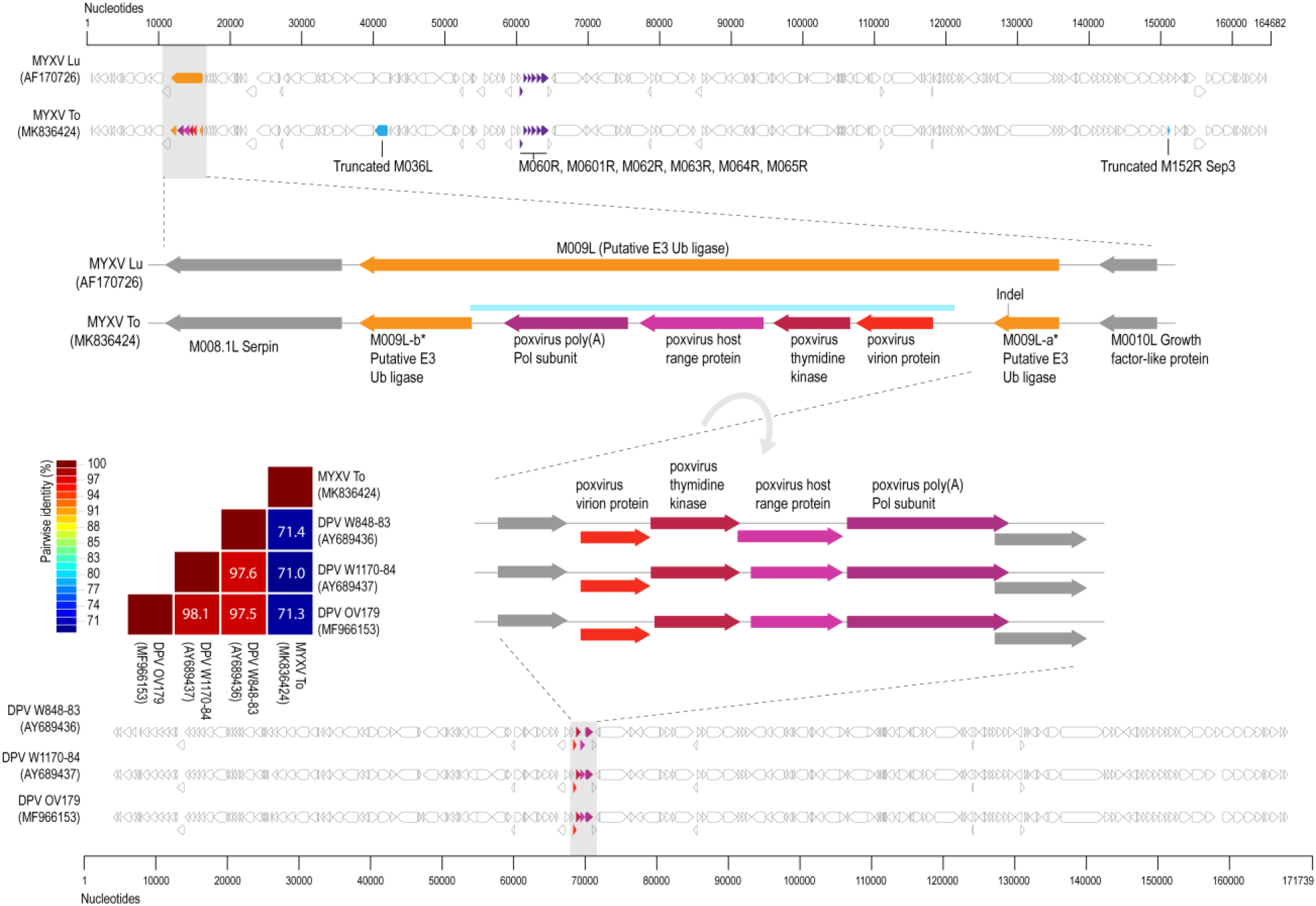
Representation of the genome organization of both MYXV-Lu (AF170726) and MYXV-To (MK836424): blue ORF illustrations represent truncated genes; purple show the location of *M060R*, *M061R*, *M062R*, *M063R*, *M064R* and *M065R* genes in both MYXV strains, orange shows the *M009L* gene (intact in MYXV-Lu and disrupted in MYXV-To) and shades of red, pink and purplerepresent the new genes inserted in MYXV-To strain after a recombinant event. The lower panel shows the only three genomes AY689436, AY689437 and MF966153 that also present a similar recombinant region. Light grey arrow indicated the inversion of the gene cassette in comparison to the three Cervidpoxvirus genomes. A pairwise identity plot with percentage pairwise nucleotide identities is provided in colored boxes for the recombinant region found in MYXV-To genome with a similar region in the three Cervidpoxvirus genomes (GenBank accession #s AY689436, AY689437 and MF966153).

### Viral genes disrupted in the new MYXV-To strain

As previously reported for MYXV isolates from feral rabbits in Australia and Great Britain, single or multiple indels that result in the disruption of ORFs are relatively common [26–28]. In the Lausanne strain, *M009L* encodes a putative E3 ubiquitin (Ub) ligase of 509 aa with a N-terminal BTB-BACK domain followed by 4 Kelch motifs [29]. Our genomic analysis reveals that ORF *M009L* of MYXV-To is disrupted by an insertion of four nucleotides (+TATA, at position 15,586-bp), causing a frameshift mutation. This indel results in a smaller truncated M009L predicted protein of 148 aa. Several reports show that this same gene is also disrupted in multiple Australian MYXV strains [28], as well as in the Californian MSW strain [16], which suggest that the disruption of this gene does not abrogate MYXV survival in the wild. Four additional nucleotides were also found in the *M036L* gene (+TTTT, position 42,007 bp), thereby creating a premature stop codon in frame within this gene. M036L is an orthologue of the O1 protein that is found in the orthopoxvirus vaccinia virus (VACV) [28]. However, the function of M036L in the MYXV virus is not reported. A previous study showed that certain MYXV field isolates carry a deletion of 89 nt in this gene [30]. However, this indel appear to have no major effects in the survival and spread of MYXV in rabbits [30]. In the MYXV-Lu strain, ORF M152R encodes a serine proteinase inhibitor (Serp3) of 266 aa [31]. In the MYXV-To strain, this gene is disrupted as a result of an insertion of a single nucleotide (+C, at position 150,688 bp), resulting in the appearance of an early stop codon. The exact biological function of Serp3 is not known in MYXV. To date, two other serpins have been identified in MYXV, Serp1 and Serp2 [32], both of which are implicated in the modulation of host inflammatory responses [33–35]. Phenotypically, the deletion of specific host range proteins inevitably results in the reduced ability of the resulting virus to infect cells or tissues of species for which the parental virus was adapted. For this reason, we consider it less likely that the truncation of *M152R* contributes to the observed virulence of MYXV-To in Iberian hares.

### Analyses of the new recombinant region of the MYXV-To strain

Analyses of the MYXV-To genome sequence revealed an insertion of ~2,800 bp in the left side of the genome (Figure 2). This new recombinant region encodes at least four genes that are predicted to encode four viral proteins that are homologous, but not identical, to the poxvirus gene families exemplified by the *M060R*, *M061R*, *M064R* and *M065R* genes from MYXV. We exploited sequence similarity searches to predict the functions of these new MYXV-To proteins. According to the obtained results, the recombinant region encodes a known virion protein (rPox-virion protein), followed by a thymidine kinase (Recombinant pox virus thymidine kinase; rPox-thymidine kinase), a C7L-like host range protein (rPox-host range protein) and a poly A polymerase subunit (rPox-poly(A) Pol subunit) (Figure 2). In the MYXV-Lu genome, the region that spans the locus at ~57,500 bp include a set of six genes that are present in all MYXV strains (*M060R* to *M065R*) [24, 36]. The predicted functions for the proteins found in the recombinant region are in accordance to those found in the ~57,500 bp region of other MYXV strains [36]. However, it should be noted that the *M062R* and the *M063R* genes that are present in all MYXV strains are not present in the new recombinant insertion region at the left end of MYXV-To (Figure 2). A BLASTn-based search for the complete recombinant region with the new four gene “cassette” revealed that this virus gene arrangement is only found in genomes (GenBank accession #s AY689436, AY689437 and MF966153) of cervidpoxviruses, for which it shares ~71% nucleotide identity (Figure 2). These results suggest that the recombinant region is derived from a new still-unreported poxvirus that shares a common ancestral origin with cervidpoxviruses. Occurrences of recombination between leporipoxviruses have been described before. In fact, it was established that the malignant rabbit fibroma virus (MRV) is a result of a recombination event between two other leporipoxviruses, the Shope rabbit fibroma virus (SFV) and MYXV [24, 25]. The recombinant MRV was capable of immunosuppression and fatal malignancy in a broader host range unlike the case of SFV but more like MYXV [26–29].

The rPox-thymidine kinase predicted protein sequence shares ~70% identity to its homologous protein from leporipoxviruses and 60-65% identity to that of capripoxviruses and cervidpoxviruses (Figure 3). The rPox-virion protein predicted protein sequence shares 73% amino acid identity to those of leporipoxviruses and 63-70% with those of centapoxviruses, capripoxviruses and orthopoxviruses (Figure 4). rPox-poly(A) Pol subunit shares the highest amino acid pairwise identity (80-86%) with those from capripoxviruses, cervidpoxviruses and leporipoxviruses (Figure 5). On the other hand, the newly identified rPox-host range protein of MYXV-To is the least conserved among the proteins found in the recombinant region, sharing only 35-40% amino acid identity with M064R protein family of centapoxviruses, cervidpoxviruses and leporipoxviruses (Figure 6). Moreover, it should be noticed that the new rPox-host range protein also shares ~40% identity to the M062R protein found in MYXV strains and RFV and ~28% amino acid pairwise identity to the M063R protein, also found in MYXV and SFV. Although the proteins found in the new recombinant region of MYXV-To share higher pairwise identity to their homologous versions found in leporipoxviruses, it should be noted that in most cases a small difference (~5% pairwise identity) segregate them from, for example, centapoxviruses and cervidpoxviruses. Moreover, and as mentioned before, the new recombinant region only presents one member of the C7L-like host range gene superfamily. In fact, leporipoxviruses constitute a unique example in the evolution of this gene family, since they encode three related C7L-like gene members in tandem, *M062R* and *M063R* and *M064R* [29]. It is suggested that the emergence of these three C7L-like gene copies in MYXV arose after two events of gene duplication [29]. In our results, we report that the new recombinant insertion region of MYXV-To only contains one predicted host range protein (Figure 2), which reinforces our hypothesis that this new gene insertion region found at the left end of the MYXV-To genome is probably not a result of a recombinant event between two leporipoxviruses, but rather between MYXV and a still-unidentified poxvirus of ungulates.

**Figure 3:**
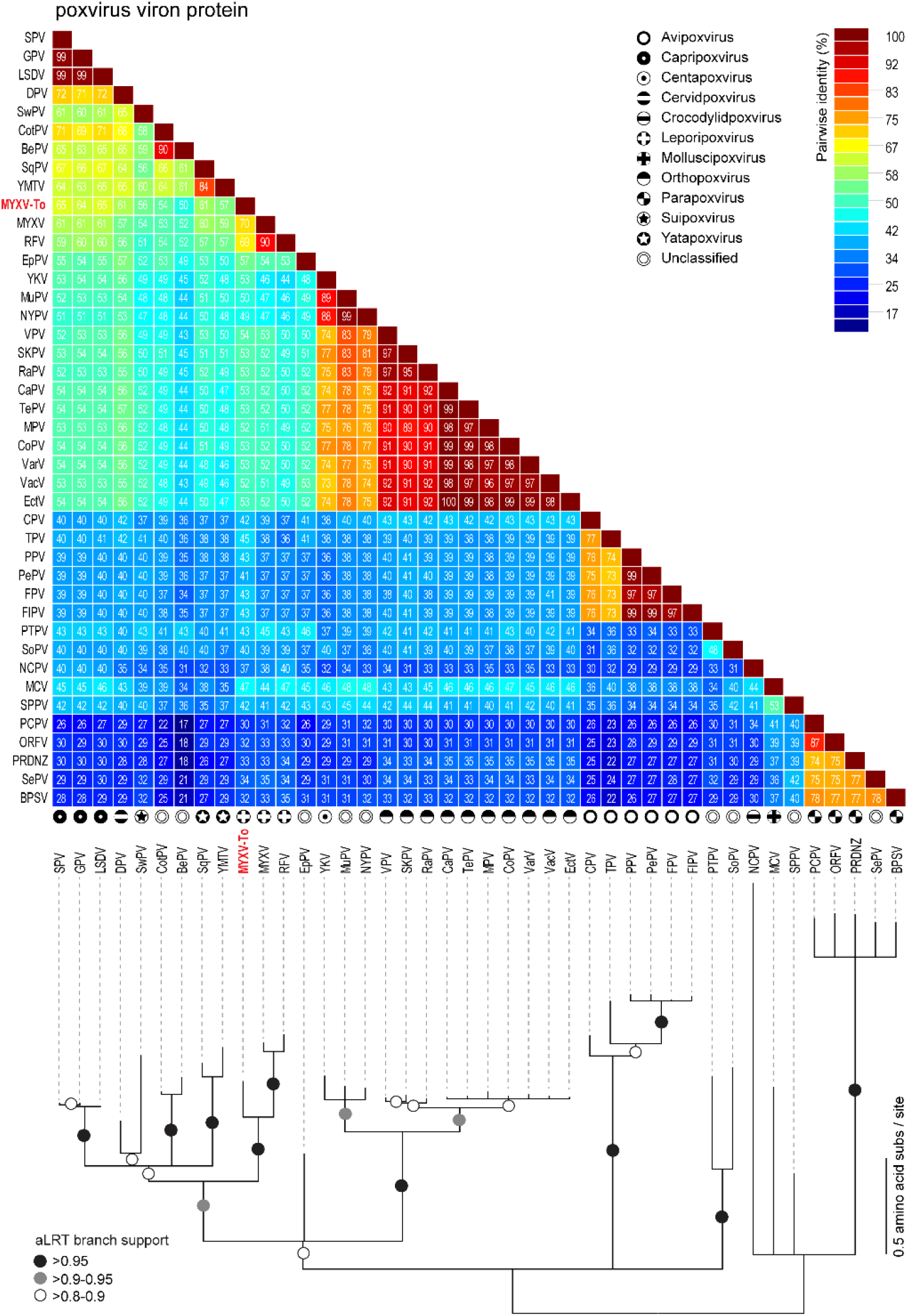
Pairwise amino acid identity matrix (upper image) and maximum-likelihood phylogenetic tree (model JTT+G) showing the relationships of the rPox-virion protein (highlighted in red) found in the recombinant region of MYXV-To strain and its homologous proteins found in representative sequences (NCBI RefSeq) of poxvirus. Branches with bootstrap support >95% are indicated with black circles whereas branches exhibiting 90%-95% and 80-90% are indicated with grey and white circles, respectively. A list of all sequences and their acronyms used for the pairwise amino acid identity matrix and phylogenetic analysis is provided in Table S1.

**Figure 4:**
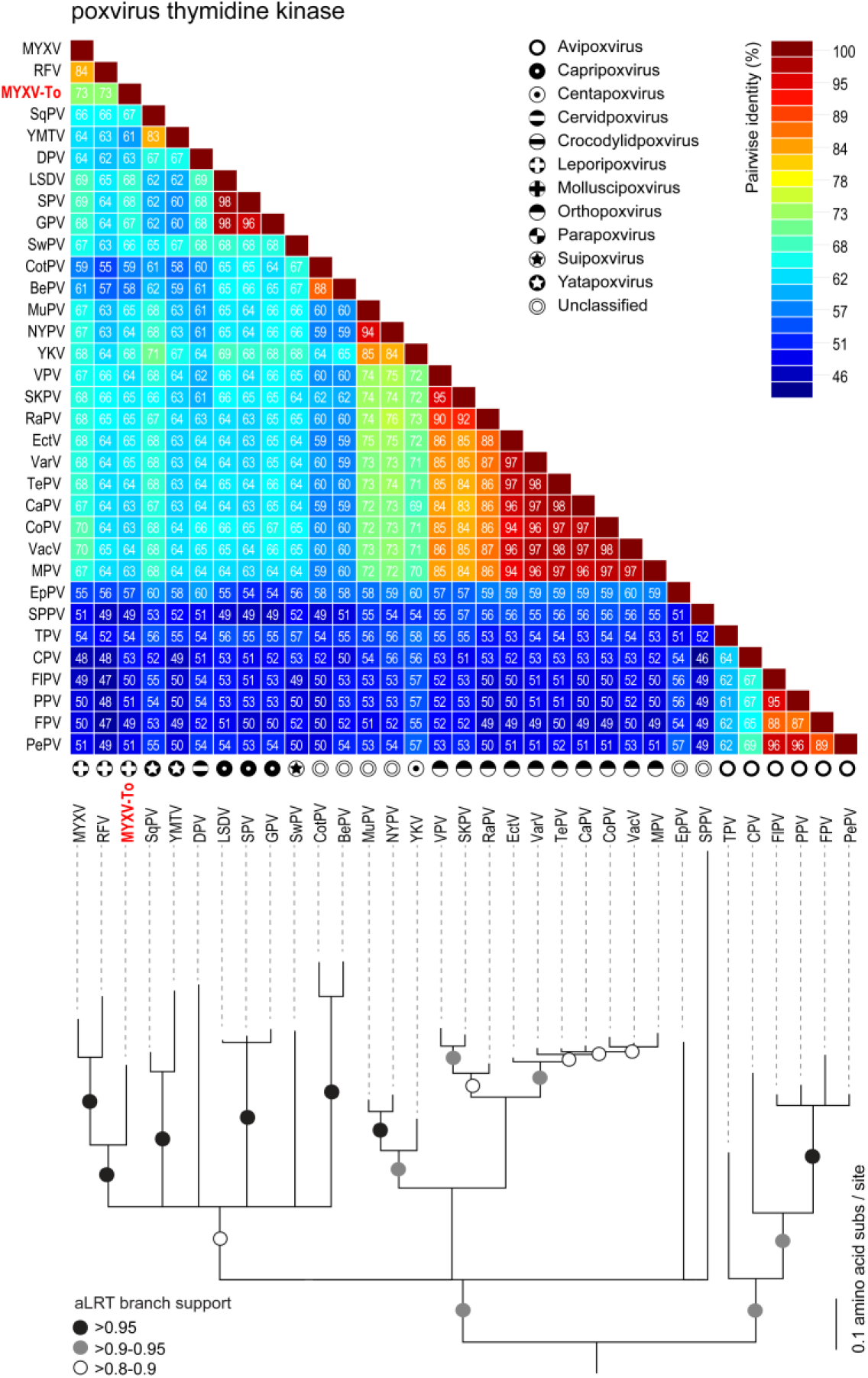
Pairwise amino acid identity matrix (upper image) and maximum-likelihood phylogenetic tree (model WAG+G+F) showing the relationships of the rPox-thymidine kinase (highlighted in red) found in the recombinant region of MYXV-To strain and its homologous proteins found in representative sequences (NCBI RefSeq) of poxvirus. Branches with bootstrap support >95% are indicated with black circles whereas branches exhibiting 90%-95% and 80-90% are indicated with grey and white circles, respectively. A list of all sequences and their acronyms used for the pairwise amino acid identity matrix and phylogenetic analysis is provided in Table S1.

**Figure 5:**
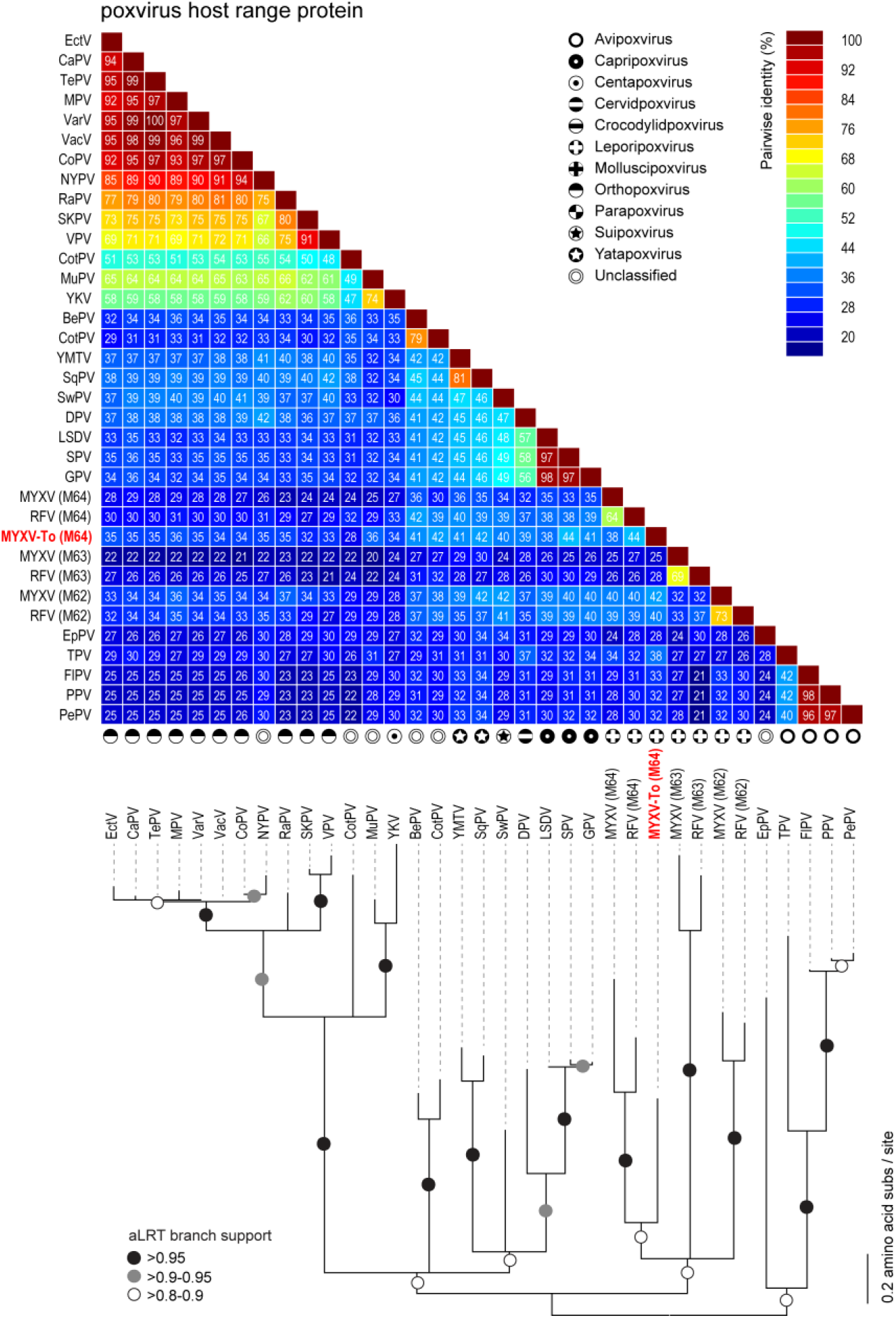
Pairwise amino acid identity matrix (upper image) and maximum-likelihood phylogenetic tree (JTT+G+F) showing the relationships of the rPox-host range protein (highlighted in red) found in the recombinant region of MYXV-To strain and its homologous proteins found in representative sequences (NCBI RefSeq) of poxvirus. Branches with bootstrap support >95% are indicated with black circles whereas branches exhibiting 90%-95% and 80-90% are indicated with grey and white circles, respectively. A list of all sequences and their acronyms used for the pairwise amino acid identity matrix and phylogenetic analysis is provided in Table S1.

**Figure 6:**
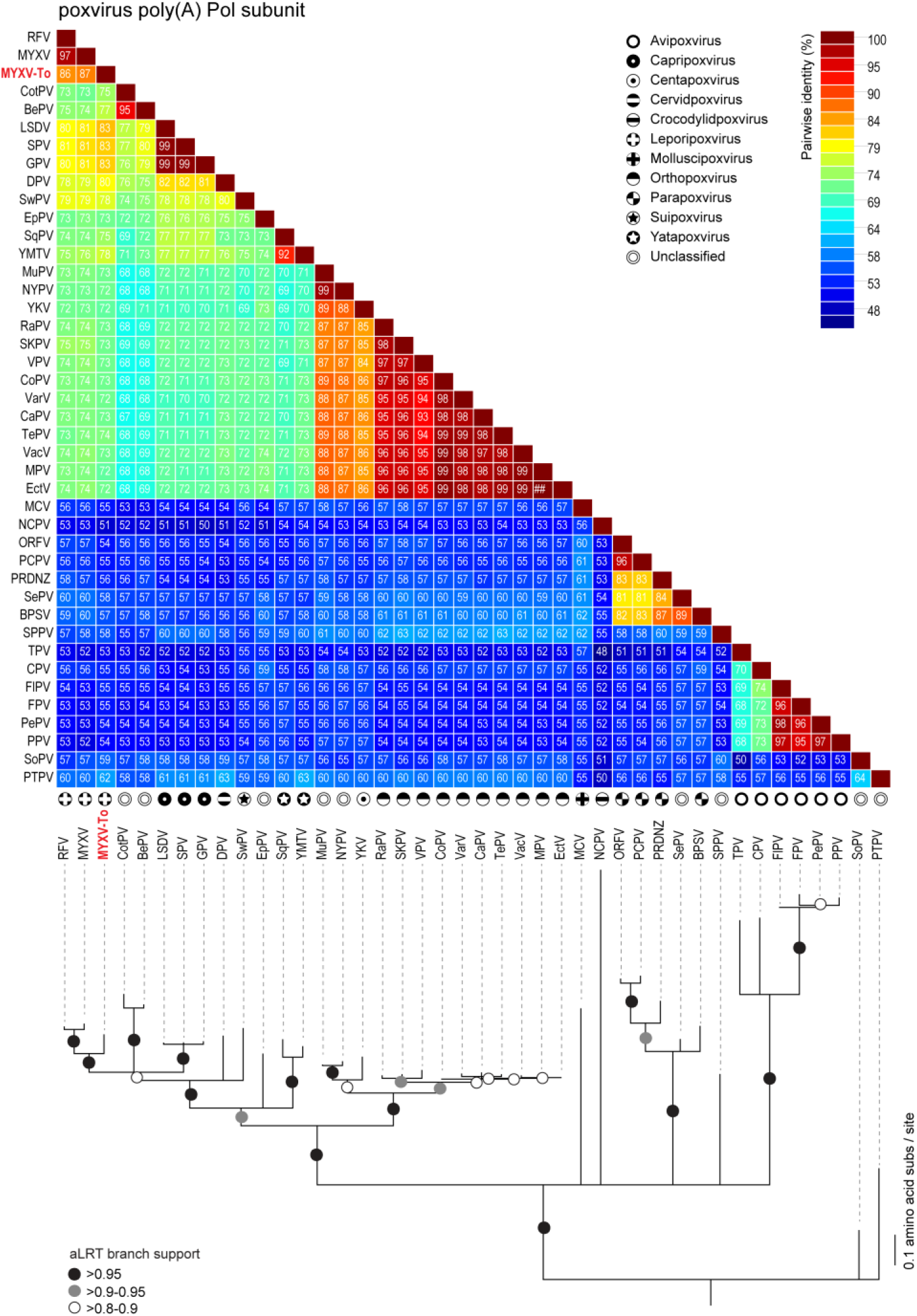
Pairwise amino acid identity matrix (upper image) and maximum-likelihood phylogenetic tree (model JTT+G+F) showing the relationships of the rPox-poly(A) Pol subunit (highlighted in red) found in the recombinant region of MYXV-To strain and its homologous proteins found in representative sequences (NCBI RefSeq) of poxvirus. Branches with bootstrap support >95% are indicated with black circles whereas branches exhibiting 90%-95% and 80-90% are indicated with grey and white circles, respectively. A list of all sequences and their acronyms used for the pairwise amino acid identity matrix and phylogenetic analysis is provided in Table S1.

### Concluding remarks

Other than the disruption of *M009L*, *M036L* and *M152R*, the MYXV-To strain has a full complement of genes present in other MYXV isolates and strains. So the question arises of how MYXV-To, with a new recombination insertion region derived from an unreported cervid-like poxvirus, became highly pathogenic in Iberian hares. Examination of the four new poxvirus genes found in the recombinant “cassette” at the left end of the MYXV-To genome, these may induce factor(s) that mediate host range and/or immunosuppression in hares, allowing the increased infection and propagation of this new virus in hares. Regarding virulence in hares, it is likely that acquisition of new genes involved in immunosuppression and/or host-range functions in specific cell types might have a preponderant role in this apparent species leaping of MYXV-To [37]. From the four new genes present in the recombinant insertion region, rPoxhost range protein is the clear candidate that suggests a possible function in novel host interactions for this new recombinant poxvirus. As mentioned before, in Lausanne strain M064R belong to the C7L-like host range factor superfamily that are known to be important for MYXV pathogenesis [38–40]. However, since this *M064R-like* gene of MYXV-To shares relatively low similarity (~40%) to its orthologous proteins found in other leporipoxviruses, it is likely that this new protein has acquired new roles, perhaps reflected by alternative host targets, compared to those in known MYXV strains. Host range proteins are defined as a group of virus-produced proteins important for the capacity of virus to infect cells or tissues of certain species [41]. The capacity of direct engagement and modulation of the host antiviral responses highlight the constant pressures exerted by the co-evolutionary arms race between host and viral pathogens [41, 42]. In fact, the high divergence observed in the new rPox-host range protein in MYXV-To might suggest that the parental virus from which this recombinant region was donated is able to replicate within a completely different host species that has a unique repertoire of anti-viral response pathways. This might ultimately result in a new poxvirus strain capable of differentially modulating the anti-viral responses of hare cells compared to MYXV, playing a critical role in species leaping and virus pathogenicity in the new host. Nevertheless, the biological implications of the new genes found in the recombination “cassette” still need to be experimentally addressed.

The data presented in this paper report that MYXV-To strain is a result of a recombinant event between a MYXV virus and a still-unreported poxvirus that shares common ancestral sequences to cervidpoxviruses. Recent reports genetically characterized a large number of European and Australian strain MYXV genomic sequences [4, 26]. Haplotypes are usually suitable for tracking the spread of MYXV virus [43]. However, the discrimination of alterations in MXYV genomes that are responsible for increased virulence grades or attenuated phenotypes is still a complicated task. While it is not yet understood what precise mechanisms allowed the MYXV-To apparently acquired virulence and species leap into Iberian hares, the genetic characterization of this novel MYXV-To strain, in combination with further studies of the proteins found in the new recombinant insertion region, will provide the foundation to a better understanding of this cross-species transmission.

**GenBank Accession #:** MK836424

## Acknowledgements

All animal sampling took place post-mortem. According to EU and National legislation (2010/63/UE Directive and Spanish Royal Decree (53/2013) and to the University of Castilla–La Mancha guidelines, no permission or consent is required to conduct the research reported herein.

FCT-Foundation for Science and Technology supported the doctoral fellowship of AAP 446 (ref. SFRH/BD/128752/2017) and the investigator grant of PJE (IF/00376/2015). This 447 article is a result of the project NORTE-01-0145-FEDER-000007, supported by Norte 448 Portugal Regional Operational Programme (NORTE2020), under the PORTUGAL 2020 449 Partnership Agreement, through the European Regional Development Fund (ERDF). This work is also supported by National Institute of Health (NIH) grants R01 A1080607.

## Author contributions

Conceptualization, PJE and AV; Methodology, AAP, AV, MB, SK and MAR; Formal Analysis, AV and AAP; Resources, PJE, CG and GM; Data Curation, AV and SK; Writing – Original Draft Preparation, AAP, ALM and PJE; Writing – Review & Editing, AV, PJE, GM and CG; Data visualization, AAP and AV; Supervision, PJE, AV and GM; Project Administration, AV and PJE; Funding Acquisition, PJE and GM.

---

**Figure S1:**
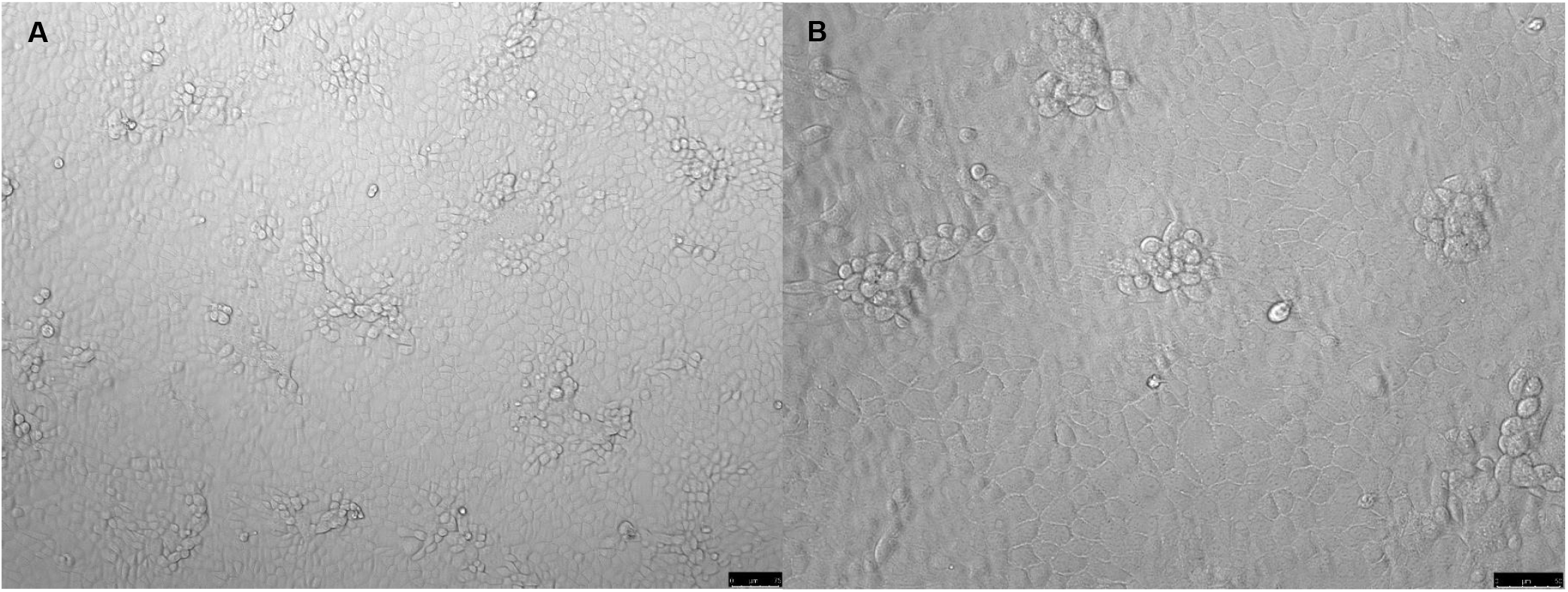
After performing a serial dilution of the purified MYXV-To virus, RK13 cells were infected and incubated for 48 hours at 37°C. At 2 days post infection, a typical MYXV cytopathic effect (foci formation) was visualized using a Leica DMI6000 B inverted microscope at 10× (A) and 20× (B).

**Figure S2:**
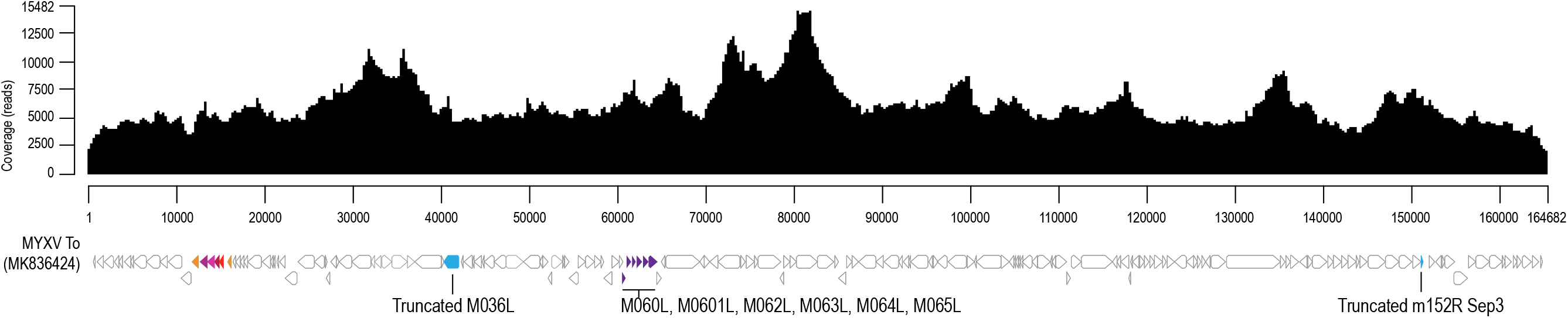
Posterior mapping of the Illumina sequencing read to the genome of MYXV-To using BBmap [16].

**Table S1:** List of representative genomes of Poxvirus (NCBI RefSeq) used for the pairwise amino acid identity matrix and phylogenetic analysis.

| Family | Subfamily | Genus | Species | Virus name(s) | Virus name abbreviation(s) | Virus isolate designation | Virus GENBANK accession | Virus REFSEQ accession |
| --- | --- | --- | --- | --- | --- | --- | --- | --- |
| Poxviridae | Chordopoxvirinae | Avipoxvirus | Canarypox virus | canarypox virus | CPV | Wheatley C93 | AY318871 | NC_005309 |
| Poxviridae | Chordopoxvirinae | Avipoxvirus | Fowlpox virus | fowlpox virus | FPV | virulent | AF198100 | NC_002188 |
| Poxviridae | Chordopoxvirinae | Avipoxvirus | Pigeonpox virus | pigeonpox virus | PPV | FeP2 | KJ801920 | NC_024447 |
| Poxviridae | Chordopoxvirinae | Avipoxvirus | Turkeypox virus | turkeypox virus | TPV | HUii24-2011 | KP728110 | NC_028238 |
| Poxviridae | Chordopoxvirinae | Capripoxvirus | Goatpox virus | goatpox virus | GPV | Pellor | AY077835 | NC_004003 |
| Poxviridae | Chordopoxvirinae | Capripoxvirus | Lumpy skin disease virus | lumpy skin disease virus | LSDV | Neethling 2490 | AF325528 | NC_003027 |
| Poxviridae | Chordopoxvirinae | Capripoxvirus | Sheeppox virus | sheeppox virus | SPV | TU-V02127 | AY077832 | NC_004002 |
| Poxviridae | Chordopoxvirinae | Centapoxvirus | Yokapox virus | yokapox virus | YKV | DakArB 4268 | HQ849551 | NC_015960 |
| Poxviridae | Chordopoxvirinae | Cervidpoxvirus | Mule deerpox virus | mule deerpox virus | DPV | W-848-83 | AY689436 | NC_006966 |
| Poxviridae | Chordopoxvirinae | Crocodylidpoxvirus | Nile crocodilepox virus | Nile crocodilepox virus | NCPV | Zimbabwe | DQ356948 | NC_008030 |
| Poxviridae | Chordopoxvirinae | Leporipoxvirus | Myxoma virus | myxoma virus | MyxV | Lausanne | AF170726 | NC_001132 |
| Poxviridae | Chordopoxvirinae | Leporipoxvirus | Rabbit fibroma virus | rabbit fibroma virus | RFV | Kasza | AF170722 | NC_001266 |
| Poxviridae | Chordopoxvirinae | Molluscipoxvirus | Molluscum contagiosum virus | Molluscum contagiosum virus | MCV | subtype 1 | U60315 | NC_001731 |
| Poxviridae | Chordopoxvirinae | Orthopoxvirus | Camelpox virus | camelpox virus | CaPV | M-96 | AF438165 | NC_003391 |
| Poxviridae | Chordopoxvirinae | Orthopoxvirus | Cowpox virus | cowpox virus | CoPV | Brighton Red | AF482758 | NC_003663 |
| Poxviridae | Chordopoxvirinae | Orthopoxvirus | Ectromelia virus | ectromelia virus | EctV | Moscow | AF012825 | NC_004105 |
| Poxviridae | Chordopoxvirinae | Orthopoxvirus | Monkeypox virus | monkeypox virus | MPV | Zaire-96-I-16 | AF380138 | NC_003310 |
| Poxviridae | Chordopoxvirinae | Orthopoxvirus | Raccoonpox virus | raccoonpox virus | RaPV | Herman | KP143769 | NC_027213 |
| Poxviridae | Chordopoxvirinae | Orthopoxvirus | Skunkpox virus | skunkpox virus | SKPV | WA | KU749310 | NC_031038 |
| Poxviridae | Chordopoxvirinae | Orthopoxvirus | Taterapox virus | taterapox virus | TePV | Dahomey 1968 | DQ437594 | NC_008291 |
| Poxviridae | Chordopoxvirinae | Orthopoxvirus | Vaccinia virus | vaccinia virus | VacV | Western Reserve | AY243312 | NC_006998 |
| Poxviridae | Chordopoxvirinae | Orthopoxvirus | Variola virus | variola virus | VarV | Ind3 | X69198 | NC_001611 |
| Poxviridae | Chordopoxvirinae | Orthopoxvirus | Volepox virus | volepox virus | VPXV | CA | KU749311 | NC_031033 |
| Poxviridae | Chordopoxvirinae | Parapoxvirus | Bovine papular stomatitis virus | bovine papular stomatitis virus | BPSV | ORFB | AY386265 | NC_005337 |
| Poxviridae | Chordopoxvirinae | Parapoxvirus | Orf virus | orf virus | ORFV | ORFD | AY386264 | NC_005336 |
| Poxviridae | Chordopoxvirinae | Parapoxvirus | Parapoxvirus of red deer in New Zealand | parapoxvirus of red deer in New Zealand | PRDNZ | HL953 | KM502564 | NC_025963 |
| Poxviridae | Chordopoxvirinae | Parapoxvirus | Pseudocowpox virus | pseudocowpox virus | PCPV | VR634 | GQ329670 | NC_013804 |
| Poxviridae | Chordopoxvirinae | Suipoxvirus | Swinepox virus | swinepox virus | SwPV | 17077-99 | AF410153 | NC_003389 |
| Poxviridae | Chordopoxvirinae |  | Tanapox virus | tanapox virus | SqPV |  | AJ293568 | NC_002642 |
| Poxviridae | Chordopoxvirinae | Yatapoxvirus | Yaba monkey tumor virus | yaba monkey tumor virus | YMTV |  | AY386371 | NC_005179 |
| Poxviridae | Chordopoxvirinae |  | Pteropox virus | pteropox virus | PTPV | Australia | KU980965 | NC_030656 |
| Poxviridae | Chordopoxvirinae |  | Squirrelpox virus | squirrelpox virus | SPPV | Red squirrel UK | HE601899 | NC_022563 |

